# Automation of citation screening in pre-clinical systematic reviews

**DOI:** 10.1101/280131

**Authors:** J. Liao, S. Ananiadou, L. G. Currie, B. E. Howard, A. Rice, S. E. Sena, J. Thomas, A. Varghese, M.R. Macleod

## Abstract

**Background:** The amount of published in vivo studies and the speed researchers are publishing them make it virtually impossible to follow the recent development in the field. Systematic review emerged as a method to summarise and analyse the studies quantitatively and critically but it is often out-of-date due to its lengthy process.

**Method:** We invited five machine learning and text-mining groups to build classifiers for identifying publications relevant to neuropathic pain (33814 training publications). We kept 1188 publications for the assessment of the performance of different classifiers. Two groups participated in the next stage: testing their algorithm on datasets labeled for psychosis (11777/2944) and datasets labeled for Vitamin D in multiple sclerosis (train/text: 2038/510).

**Result:** The performances (sensitive/specificity) of the most promising classifier built for neuropathic pain are: 95%/84%. The performance for psychosis and Vitamin D in multiple sclerosis datasets are 95%/73% and 100%/45%.

**Conclusions:** Machine learning can significantly reduce the irrelevant publications in a systematic review, and save the scientists’ time and money. Classifier algorithms built for one dataset can be reapplied on another dataset in different field. We are building a machine learning service at the back of Systematic Review & Meta-analysis Facility (SyRF).

## 1. Introduction

An increasing amount of *in vivo* research is published every day and as a result it has become virtually impossible for researchers to keep up-to-date with the progress in their field. Emerging findings have conventionally been interpreted and synthesised in review articles, but these often cite selectively from the literature, may be written with a specific purpose such as supporting a grant application, and may be partisan. Systematic review, which often but do not always includes a metaanalysis, is an alternative approach that seeks to identify and consider all relevant available evidence to provide an unbiased summary of existing research. This might be used for instance to identify gaps in research knowledge, or to establish whether there is sufficient evidence to proceed for instance to clinical trial of a novel intervention; and allows both an assessment of risks of bias in the research literature and summary estimates of observed biological effects. While this approach provides powerful information, it is a very time-consuming process and a systematic review is often outdate by the time it is finished and reach its publication stage.

Rapid developments in text-mining and machine learning techniques and their successful applications in various areas encouraged us, as practitioners of systematic reviews, to explore the possibility that these techniques might be used to accelerate the process of systematic review and meta-analysis. Even more exciting, it hints a possible future with a “living” systematic review machine – a process through which a systematic review might be updated automatically to include consideration of newly published research.^2^. Updating preclinical systematic reviews in an automated fashion would make such systematic reviews current and relevant, and reduce the burden of conducting such reviews would enable their more widespread adoption.

Text-mining has previously been tested, used and analysed in clinical systematic review area [4]. However, because experimental designs, epistemological approaches and manuscript styles are very different for in vivo research compared with human clinical trials, techniques successful in systematic reviews of clinical trial data may not be as useful in systematic reviews of preclinical studies. Since preclinical research makes up around three quarters of biomedical research expenditure, it is important that tools to summarise the research which results are available.

There are several stages in a systematic review that are time consuming. Here we focus on one of the early and more mechanical stages: reference selection. During reference selection researchers read through all references acquired from searches of online databases and decided whether each reference is relevant to the project, making that judgement against a number of predefined inclusion and exclusion criteria. Depending on how broad the scientific questions are, the number of references to screen varies but may reach several hundred thousand.

We invited five text-mining groups to apply their text mining techniques to sets of screened search results that we had collated in the context of our previous systematic reviews. We received 13 classifiers that used different combination of nature language processing settings, feature selection methods, and classification algorithms. A study protocol was developed to set out our approach, and to allow the reader to have confidence that our analysis approach was determined in advance of data collection.

## 2. Data

To understand the feasibility of the machine learning approach to reference selection across a range of situations we used three different datasets from different pre-clinical research fields.

### 2.1 Neuropathic pain dataset

The largest dataset is from a systematic review and meta-analysis study of neuropathic pain [5]. We carried out a search of 5 online databases for studies reporting animal experiments modelling neuropathic pain where a behavioural outcome was reported. This search, carried out in September 2012, identified 33,814 unique publications. Two independent investigators screened the title and abstract of these publications against pre-defined inclusion/exclusion criteria. They agreed on inclusion or exclusion for 33,174 publications, and after consultation with a third reviewer a decision was reached on all 33,814 publications, 6506 of which were included. These publications form the dataset for this study and are now being included in a family of systematic reviews categorised by the pain model used. The dataset of 33,814 publications with known status for “included” or “excluded” was sent to five machine-learning groups to train the classifiers.

As stated in the protocol [6], in November 2015 we repeated the search using the same search terms and retrieved 11880 new publications. We randomly selected 1188 (10%) of these and two independent reviewers reviewed them against the inclusion/exclusion criteria of the neuropathic pain project. All publications in the updated search were sent to the text-mining groups for classification. The information provided to the classifiers consisted of title, author, abstract, year, and classification label if available. The performance of the classifiers is assessed against the human decision of the 1188 reviewed publications.

### 2.2 Psychosis dataset

Text-mining algorithm perform differently depending on different complexity of the dataset and inclusion/exclusion criteria. To test the generality of the built algorithms and to learn about response of classifier parameters for different dataset, we send out another dataset - publications screened against animal model of psychosis. Because of the complexity of the animal models of psychosis, reviewers have screened this dataset against more complicated inclusion/exclusion criteria, and as a result included broad range of animal models. The high complexity of the labelling logic provides a new challenge for the classifiers, which is optimised for the neuropathic pain dataset. This test provides an insight of the reusability of a text-mining classifier. While the pipelines of the classifiers do not change, all hyper-parameters and parameters are optimised during training the classifiers with the new dataset.

In this dataset, we carried out a search on PubMed for studies reporting animal models of psychosis reporting a behavioural outcome. This search, carried out in January 2014, identified 14,721 unique publications. Two independent researchers screened these publications against predefined inclusion/exclusion criteria. They agreed on inclusion or exclusion for 13,086 publications, and after consultation with a third reviewer a decision was reached on all 14,721 publications, 4,335 of which were included. These are now being included in a family of systematic reviews categorised by the psychosis model used. We sent out a random sample of 11,777 records (80% of the dataset) for training purpose and reserved 2,944 (20%) for testing purpose.

### 2.3 Vitamin D in multiple sclerosis (MS) dataset

Another aspect of the dataset known to impact the performance of classifier is the prevalence of the positive label. When the prevalence is low, a small number of false negative decisions could significantly harm the sensitivity (recall) of the classifier. The third dataset we sent to our text-mining groups is the publications labelled against inclusion/exclusion criteria of the human and animal evidence of Vitamin D in multiple sclerosis. As with the other two datasets, this reference selection dataset is also created by three independent reviewers. The inclusion/exclusion criteria are less complex in this project as only one treatment is included. However, only 83 out of 2548 references were included. By training and testing the classifiers using this low prevalence (3%) dataset, we can study the performance dependence on prevalence of a text-mining classifier optimised for another dataset. We sent out a random sample of 2038 records (80% of the dataset) and withheld 510 (20%) for testing purpose.

Table 1 summarizes the level of complexity, the number of references included in training and testing three datasets and the prevalence of each datasets.

**Table 1.** Summary of the datasets.

|  | Neuropathic Pain | Psychosis | Vitamin D in MS |
| --- | --- | --- | --- |
| **Complexity of Inclusion Criteria** | Medium | High | Low |
| **Training Dataset Size** | 33814 | 11777 | 2038 |
| **Training Dataset Prevalence** | 19% | 29% | 3% |
| **Testing Dataset Size** | 1188 | 2944 | 510 |
| **Testing Dataset Prevalence** | 30% | 29% | 3% |

All three datasets are from completed screening projects. Due to the nature of the three projects, the model complexities are different and the original search strings have unavoidable differences in their sensitivity and specificity. Datasets of neuropathic pain and psychosis contains more complex disease model induction and outcome assessments and hence more complex inclusion criteria. The Vitamin D in multiple sclerosis (MS) project only investigated one intervention, so it contains a much simpler inclusion criteria but also has a much lower prevalence. We will discuss the impact of those differences.

## 3. Methods

### 3.1 Building Classifiers using Neuropathic pain dataset

Five text-mining groups participate in this project and built literature selecting classifiers for neuropathic pain dataset: Institution of Education (IOE), London Uk; National Centre forText Mining (NaCTeM), Manchester UK; SWIFT [8], DoCTER [7] and our in house CAMARADES group at the University of Edinburgh [9].

#### 3.1.1 Core algorithm and feature selection settings

There are many different machine-learning algorithms and techniques that may be used in building a good classifier. To gain insights into the classifier built by different machine learning groups, we asked them to provide the classification algorithm, feature settings, and feature preparation procedures for each submitted result.

For the main study using neuropathic pain dataset, we receive results from 13 classifiers produced by 5 text-mining groups. Table 2 lists the details of each algorithm. We report the details we received without modification. The sequence of the classifiers represents the sequence in which the results were received. The numbers of classifiers submitted are different for different groups, and some groups submitted classification results at different time points.

**Table 2.** Details of the algorithms and main settings of 13 received classifiers. Some of the information was not available or was not provided

| Classifier | Prioritisation | Seed Size | Classifier | Feature Selection | Feature No. | Kernel | Other Settings |
| --- | --- | --- | --- | --- | --- | --- | --- |
| 1 | N | 33814 | SVM | BOW | / | Linear | / |
| 2 | Y | 33814 | SGD | BOW | / | Linear | / |
| 3 | N | 33814 | SVM | LDA | / | Linear | / |
| 4 | N | 33814 | Ensemble of NB, SVM, kNN, and LDA | BOW | / | Linear | / |
| 5 | Y | 33814 | SGD | BOW | / | Linear | / |
| 6 | N | 33814 | SVM | BOW | / | Linear | / |
| 7 | N | 4058 | SVM | BOW | 207 | Radial | C = 0.05, Gama = 0.008 |
| 8 | N | 33814 | Ensemble of NB, SVM, kNN, and LDA | BOW | / | Linear | / |
| 9 | N | 20228 | SVM | BOW | 16404 | Radial | C=2, Gama = 0.002 |
| 10 | Y | 33084* | Ensemble of multiple classifier types | LDA & BOW | 16404 | / | Target 99% sensitivity |
| 11 | Y | 33084* | Ensemble of multiple classifier types | LDA & BOW | 16404 | / | Target 95% sensitivity |
| 12 | Y | 33084* | Ensemble of multiple classifier types | LDA & BOW | 16404 | / | Target 85% sensitivity |
| 13 | Y | 33084* | Ensemble of multiple classifier types | LDA & BOW | 16404 | / | Target 80% sensitivity |

As shown in table 2, three different machine learning classification algorithms are used: Support vector machine (SVM), stochastic gradient descent (SGD) and ensemble of multiple classifier types, in various combinations with two different feature selection methods, Bag of Words (BOW) and Latent Dirichlet allocation (LDA). Prioritisation is used in half of the classifiers. Most classifiers use linear kernel. SVM and SGD with linear kernel are thought to be among the best approaches for classification jobs (ref), while BOW and LDA are still competing for recognition as the best approach for the feature selection method.

#### 3.1.2 Text preparation

Optimization of text preparation is important. As the classification algorithms become more and more standard, the procedures of text preparation before training, e.g. Lemmatisation and punctuation removal, and the sequence in which these are conducted can be key in differentiating a good classifier from the best. To learn more about the importance of text-preparation we asked each text-mining group to provide their text preparation procedures and the sequence, and we list these in table 3.

**Table 3.** Feature selection and processing details

| Classifier | To lower | Remove new lines | Remove numbers | Remove punctuations | Remove stop words | Stemming / Lemmatisation | Strip whitespace | Tokenizer (Details) | Sentence splitter |
| --- | --- | --- | --- | --- | --- | --- | --- | --- | --- |
| 1 | 1 | Yes | 5 | 4 | 2 | 3 | Yes | Yes | Yes |
| 2 | 1 | No | 2 | No | 3 | No | 4 | / | / |
| 3 | 1 | Yes | 5 | 4 | 2 | 3 | Yes | Yes | Yes |
| 4 | Yes | No | No | Yes | Yes | No | Yes | Yes | No |
| 5 | 1 | No | 2 | No | 3 | No | 4 | / | / |
| 6 | 1 | Yes | 5 | 4 | 2 | 3 | Yes | Yes | Yes |
| 7 | 1 | 2 | 3 | 4 | 5 | 6 | 7 | No | No |
| 8 | Yes | No | No | Yes | Yes | No | Yes | Yes | No |
| 9 | 1 | 2 | 3 | 4 | 5 | 6 | 7 | No | No |
| 10 | Yes | Yes | No | Yes | Yes | Snowball (English) | Yes | Whitespace tokenizer (after punctuation removal) | No |
| 11 | Yes | Yes | No | Yes | Yes | Snowball (English) | Yes | Whitespace tokenizer (after punctuation removal) | No |
| 12 | Yes | Yes | No | Yes | Yes | Snowball (English) | Yes | Whitespace tokenizer (after punctuation removal) | No |
| 13 | Yes | Yes | No | Yes | Yes | Snowball (English) | Yes | Whitespace tokenizer (after punctuation removal) | No |

Where the sequence is not known we simply list whether the general approach was used or not. Some groups provide more details for specific procedures. Procedures that were done by some included: lower the cases, remove stop words, and strip white spaces. Procedures not done included: remove numbers, remove new lines, remove punctuations, stemming/lemmatisation, tokenizer, and sentence split.

### 3.2 Testing built classifiers on Psychosis and Vitamin D Dataset

To understand the possibility to use an classifier built for one dataset, we requested our text-mining groups to apply their classifier built for the neuropathic pain dataset to the psychosis and Vitamin D datasets. IOE and NaCTeM participated in this stage of study, and their results are shown in section 4.

## 4. Result

For most of the pre-clinical research, the number of retrieved literatures for search engines, e.g. PubMed, is often very large. A typical prevalence of the relevant publications from this pool ranges from 3% to 20%. Hence, for machine-learning application in reference screening stage of pre-clinical systematic review, sensitivity (recall) is the satisfactory specification metric while the specificity is the optimising metric.

As there was no similar study before for pre-clinical research, and the environment is highly different for pre-clinical and clinical systematic review, we did not have an evidence-based expectation value of sensitivity for this study. We encourage our machine learning collaborators provide what they thought was the best results. As the believed Bayes error rate of dual manual reference screening at title and abstract stage is at 2%, we aspire to a sensitivity of 95%, and indeed this figure was later endorsed as a reasonable target in by an expert panel convened by the SLiM consortium in London in late 2017. However, this number is not considered as an absolute cut-off line.

Because each systematic review is built around a unique scientific question and its search result, desirable specification metrics (and in particular the consequences of not identifying 1%, or 5%, or 10% of a literature) could be different for different systematic reviews. Judgement of the optimal approach will of course include computation time and training data required. We will further discuss the generality of the machine-learning algorithms in the discussion session.

We calculate the confusion matrix and list the sensitivity, specificity, precision, F1 score, utility, and utility (PPV). We define utility and Utility (PPV) as following:

### 4.1 Neuropathic pain dataset

We received 13 classifiers for the study using neuropathic pain data set, and tested them against our reserved test dataset and list their performance in table 3. Two of the 13 classifiers are derived from raw scores provided by machine-learning teams, who sent out scores rather than classifications by optimising metrics. In these cases we chose one cut point on the score that secured a sensitivity of 95% and another cut point that secured specificity at 75%.

Four of the classifiers have sensitivity above 95%. Among those four classifiers, classifier #3 has the highest specificity at 84%. Classifier #11 with sensitivity at 87% and specificity at 95% is likely to filter out more irrelevant studies at the cost of missing some relevant studies. There is a trade-off relation between the satisfactory and optimisation specification metrics, i.e. sensitivity and specificity, which provides an opportunity to systematic reviewers to choose the best pair of metrics for each projects.

### 4.2 Psychosis dataset

Two machine-learning teams (IOE and NaCTeM) participated in the following up study and provided results for psychosis data; there were four sets of results. The algorithm/feature settings are the same as those built for neuropathic pain study. We reserved 20% of data for testing and the performances are listed in table 4.

**Table 4.** Classifier performances for neuropathic pain dataset

| Classifier | Sensitivity | Specificity | Precision | F1 Score | Utility | Utility (PPV) |
| --- | --- | --- | --- | --- | --- | --- |
| 1 | 0.97 | 0.73 | 0.94 | 0.75 | 1.01 | 1 |
| 2 | 0.97 | 0.67 | 0.97 | 0.71 | 1 | 1 |
| 3 | 0.95 | 0.84 | 0.98 | 0.82 | 0.99 | 0.99 |
| 4 | 0.93 | 0.48 | 0.94 | 0.59 | 0.96 | 0.96 |
| 5 | 0.91 | 0.87 | 0.87 | 0.82 | 0.96 | 0.95 |
| 6 | 0.87 | 0.95 | 0.98 | 0.87 | 0.92 | 0.92 |
| 7 | 0.81 | 0.52 | 0.96 | 0.56 | 0.85 | 0.84 |
| 8 | 0.80 | 0.6 | 0.87 | 0.59 | 0.84 | 0.83 |
| 9 | 0.74 | 0.92 | 0.89 | 0.77 | 0.8 | 0.8 |
| 10 | 0.98 | 0.71 | 0.99 | 0.74 | 1.01 | 1.01 |
| 11 | 0.90 | 0.92 | 0.95 | 0.86 | 0.95 | 0.94 |
| 12 | 0.78 | 0.97 | 0.91 | 0.84 | 0.84 | 0.84 |
| 13 | 0.71 | 0.98 | 0.89 | 0.82 | 0.78 | 0.78 |

Utility = $((19 \times \text{specificity}) + (1 - \text{sensitivity})) / (1 + 19)$ , Utility (PPV) = $((19 \times \text{precision}) + (1 - \text{sensitivity})) / (1 + 19)$

We learnt from the results of neuropathic pain dataset and set up the satisfactory sensitivity at 95%. Classifier #4 provides the best specificity. The differences between the lowest and highest specificity is only 5%. While pre-clinical models and treatments of psychosis have higher variety and more complicated in description, and the classifiers are optimised for neuropathic pain dataset, the specificity remained around 70%. It is worth noticing that the number of training set is at best less than one third of that available for the neuropathic pain dataset. We will discuss the impact of training dataset on the classifier performance in the discussion session.

### 4.3 Vitamin D in MS dataset

We sent out dataset screened for vitamin D in MS to our machine-learning teams. IOE and NaCTeM provided four classification results in total. The most striking characteristic of this dataset is the low prevalence (3%). We retained 20% of data to test the performance as listed in table 5.

**Table 5.** Classifier performances for Psychosis dataset

| Classifier | Sensitivity | Specificity | Precision | F1 Score | Utility | Utility (PPV) |
| --- | --- | --- | --- | --- | --- | --- |
| 1 | 0.95 | 0.67 | 0.55 | 0.7 | 0.99 | 0.98 |
| 2 | 0.95 | 0.69 | 0.57 | 0.71 | 0.99 | 0.98 |
| 3 | 0.95 | 0.72 | 0.59 | 0.73 | 0.99 | 0.98 |
| 4 | 0.95 | 0.73 | 0.60 | 0.74 | 0.99 | 0.98 |

**Table 6.** Classifier performances for Vitamin D in MS dataset

| Classifier | Sensitivity | Specificity | Specificity | F1 Score | Utility | Utility (PPV) |
| --- | --- | --- | --- | --- | --- | --- |
| 1 | 1.00 | 0.36 | 0.04 | 0.09 | 1.02 | 1.0 |
| 2 | 0.93 | 0.58 | 0.06 | 0.12 | 0.97 | 0.94 |
| 3 | 1.00 | 0.45 | 0.05 | 0.10 | 1.02 | 1.0 |
| 4 | 0.93 | 0.84 | 0.15 | 0.26 | 0.98 | 0.94 |

While we still want to keep the sensitivity at 95%, we notice that the specificity significantly dropped. The low prevalence of positive instances and the limited number of studies in this dataset lead to the discrete results of the sensitivity. At sensitivity of 100%, the highest specificity we have is at 45%, which is much lower comparing to the previous two studies. Classifier #4 provides an alternative optimised result: at 93% sensitivity, the specificity can reach 84%. In conclusion, the combination of low prevalence coupled with a small training dataset strongly compromises the performance of machine-learning classifiers.

## 5. Discussion

### 5.1 Core Algorithm

#### 5.1.1 Discussion for different classifiers

Three algorithms are used in all three studies: SVM, SGD and ensemble of multiple classifier types. There is no clear difference in performance among three algorithms. For those building textmining classifiers for relatively complex concepts, either SVM or SGD appear to be a good choice for core algorithm to start with.

#### 5.1.2 Execution time

There are two considerations of execution time of importance for this kind of classifier. The first is training time, which is the time need to train the machine to build the classifier, including the time to prepare the features from the datasets and then to train the classifier. While the time required depends on the algorithms and the computation power of the server, training a dataset of 30,000 references with title and abstract takes several hours, but not days.

Once the classifier is trained, it is ready to be used on unclassified data. The 2^nd^ execution time is the application time of the classifier on the unclassified data. This time is much shorter and is usually around 30 minutes for 10,000 references. These time references are based on optimised machine learning algorithm performance running on a server in NaCTeM.

#### 5.1.3 Training data

The main purpose of building an automatic classifier to perform the reference selection is to save systematic reviewers’ precious research time. One question we also get from researchers who want to use machine learning to help with reference selection is ‘how much do I need to screen to training the machine?’ To answer this question, we plot the learning curve of a classifier, i.e. the graph of the performance against the amount of training data used, as shown in figure 1.

**Figure 1.**
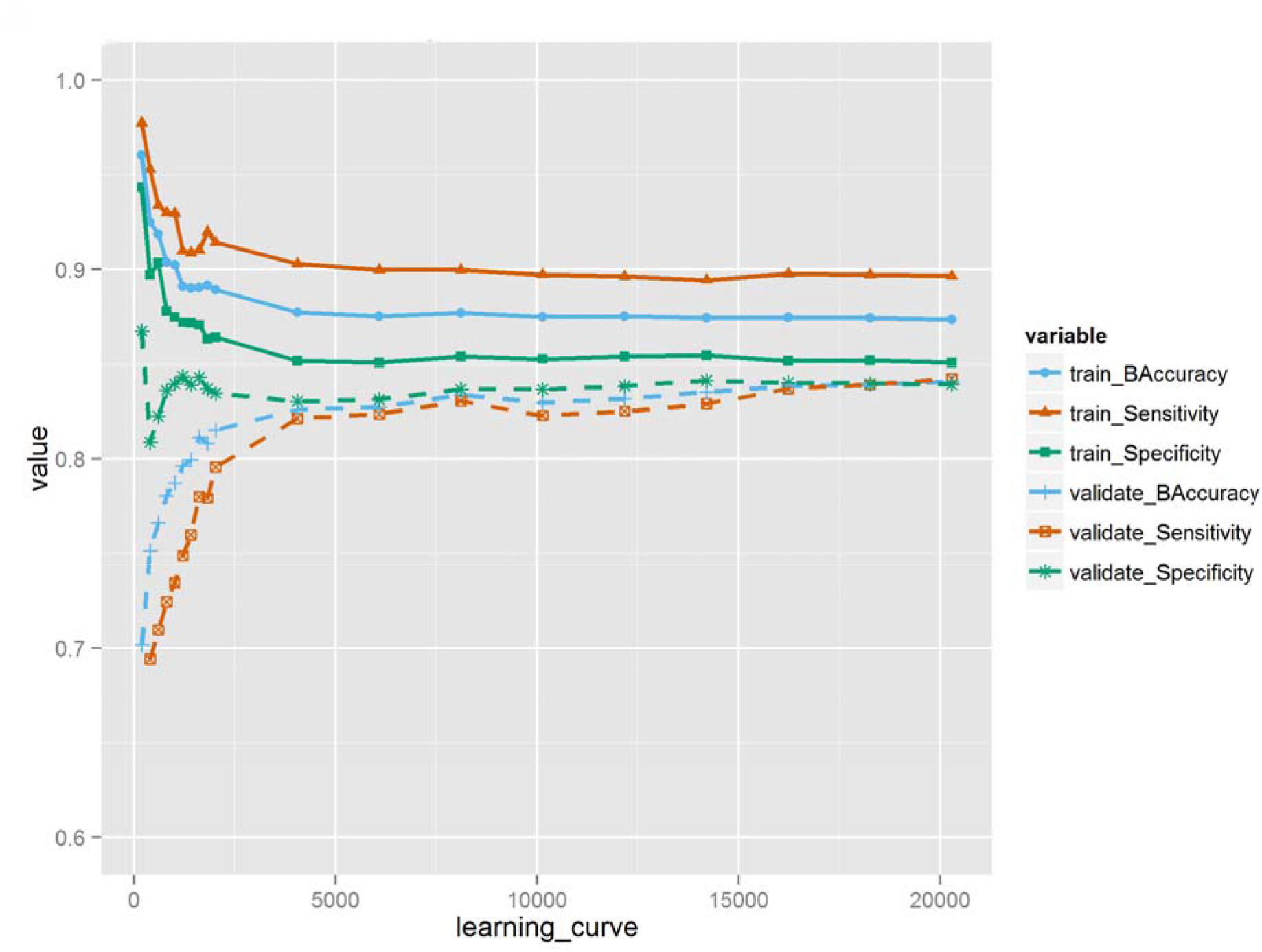
Classifier performances for Psychosis dataset

Figure 1 shows the learning curve for classifier #8 during training data using neuropathic pain dataset. There is a clear positive correlation between the size of training dataset and the performance on the validation sets while the improvement of performance slows down as the training data size increases significantly. There are two critical numbers worth noticing. The performances of the machine built with less than 2000 training data are low and unstable, which means, for neuropathic pain project, it only makes sense to use machine learning to aid reference selection if the total number of publications obtained from the search engine is larger than 2000. When using 2000 training data, the performance of the classifier is relatively stable but has a much lower sensitivity and specificity. When the size of the dataset reaches 5000, the performance increases to near the optimised point and after that, the slope of the improvement is much slower. This is the 2^nd^ critical number: the number when the improvement of the classifier reaches a plateau, and after that it will much more effort to improve the classifier performance by increasing the dataset.

While critical numbers of training datasets highly depend on the complexity of the inclusion criteria and the performance demanded, we hope these numbers can guide researchers in selecting useful machine-learning methods.

#### 5.2 Choice of cut-off line in experiment and practical use

In this study multiple groups provides scores for each publication (usually a number between 0 and 1) instead of the traditional classification result of 1 (“include”) or 0 (“exclude). We see this as desirable, as it allows the user to have flexibility in establishing, essentially, what type of mistake they would prefer the machine to make. Ideally, a well-trained classifier should have a distribution of score as shown in figure 2, a normal distribution with a tail bump. Figure 2 plots the histogram of the scores from the training results of IOE.

**Figure 2.**
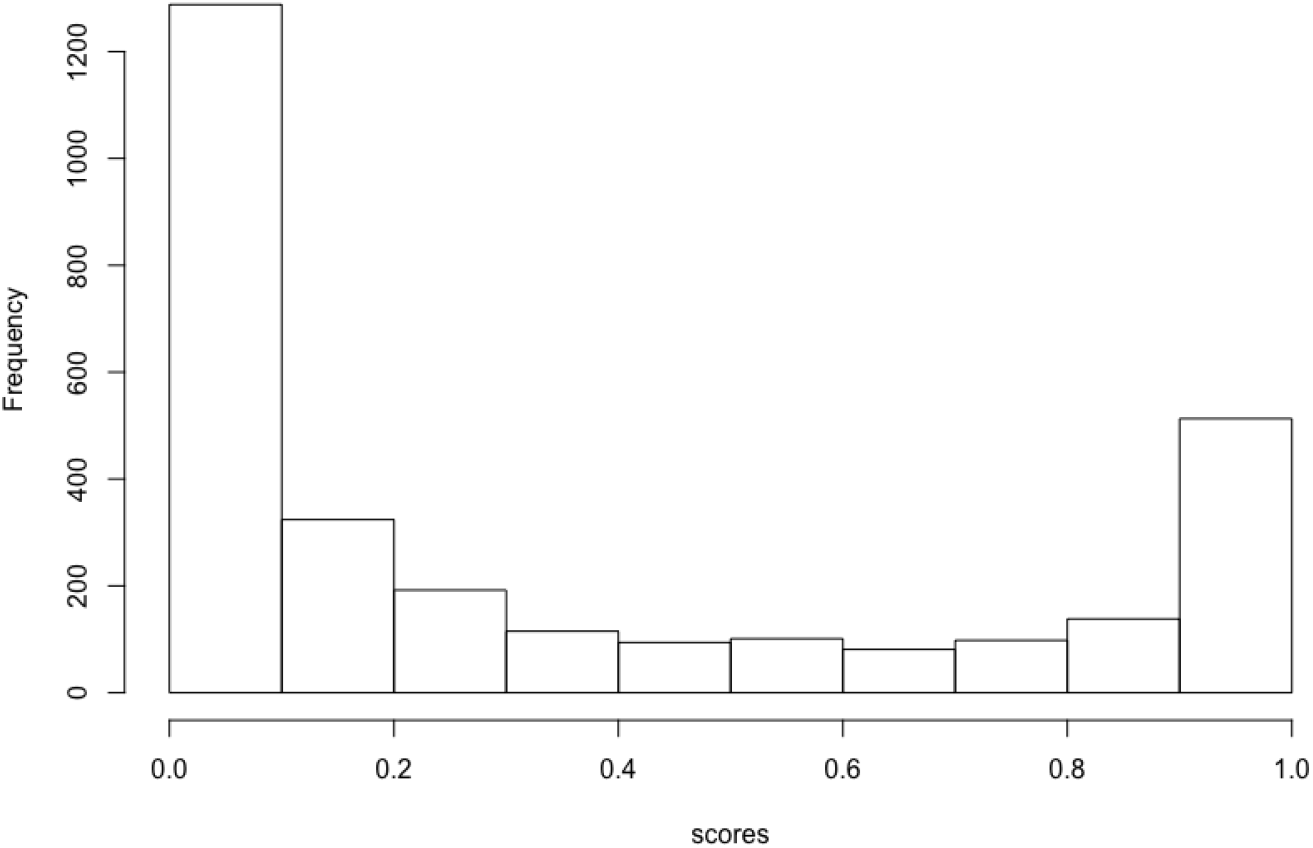
Classifier performances for Psychosis dataset

The ideal cut-off for this classifier is around 0.8. Again, depending on the complexity of the dataset and requirement of the project, the ideal cut-off line varies, but with access to the score the systematic reviewer can make the choice of the trade-off point between sensitivity and specificity after obtaining the results for performance in the validation set by modifying the cut-off line dividing inclusion and exclusion.

#### 5.3 Contribution of different feature types

Bag of words and latent Dirichlet allocation were the most common feature types used. BOW is the classical way of mapping documents into the space of words while each word is treated as an arbitrary discrete atomic symbol. It is intuitive, the feature building process takes less time, and there are fewer parameters to tune. However, BOW naturally lacks of information regarding the relationships between the individual words, and it leads to large data sparsity, which then means the model is more data thirsty and takes more time to train. Latent Dirichlet allocation is a popular alternative feature type used in text mining. It maps documents into groups of topics instead of words, and the topics are learned using all the provided documents (including unlabelled documents). Compared to BOW, LDA takes longer time to complete at the feature preparation stage but uses less time in training the classifier. As for the performance of the classifier, by combining table 2 and table 4, we concluded that there is no obvious difference of performance between classifiers using BOW and LDA. However, there is a drawback of LDA: because LDA uses all documents to build the topic models, the classifier has to be retrained whenever there are new documents.

#### 5.4 Contribution of different feature preparations

Increasing experience with text mining approaches suggests that feature preparation could make a significant impact on the final performance of the classifier. By combining information from table 3 and table 4, we learn that the most necessary steps are lowering the case of the words, removing stop words, and stripping white spaces. Recommended steps are removing new line, remove punctuations, stemming/Lemmatisation, tokenizer, and sentence splitter.

#### 5.5 Updating an systematic review and concept drift

Because of the rapid publishing speed of pre-clinical studies, updating systematic reviews to ensure they provide contemporary information is increasingly important. For systematic reviews with large datasets, one may build a classifier using machine learning and apply it to the references in the updated search (as we have done here with neuropathic pain). There is however a potential complication that may compromise the performance of the classifier on the new sets of references. Concept drift occurs when the terminology used in a literature changes over time; in pre-clinical systematic review it implicates the changes of the way one describe a disease model or new treatment method that did not exist in the earlier sets of references. The evolution of the search engines that include different set of publications at different times may also have an impact. The observation that the prevalence of included citations in the neuropathic pain dataset differs between the original search and the updated search is possibly due to the changes in both online search engines and their embedded heuristics as well as concept drift. While there is no way to measure the impact at this moment, we are aware of concept drift and its possible impact on the performance of the classifier. We believe that testing the performance on updated searches is essential to ensure accuracy, and updating the training data with newly manually screened results might help reduce the impact.

### 6. Conclusion

We performed a pilot study of accessing machine-learning applications in pre-clinical systematic review using three different dataset and collaborating with five different machinelearning groups. We have found that the performance of the automated classifiers is pronounced for the original neuropathic pain dataset, with multiple classifiers reaching sensitivity at ~95% and specificity > 80%. Some of the classifiers are tested against two different dataset with different complexity of the inclusion criteria, and the results are promising, which increases our confidence in the generality of the classifier. The test of the classifier on a dataset with different prevalence shows that low prevalence coupling with low number of total references can limit the performance of the classifier.

A further investigation of the learning curve of the classifier shows that systematic review projects with more than 2000 retrieved studies should consider using machine-learning approaches to help with the reference selection, while keep in mind that it may require as many as 5000 labelled publications to train the classifier to a satisfying performance.

To help researchers to perform systematic review using machine learning without their needing to understand and implementing machine-learning algorithms, we have collaborated with NaCTeM and IOE to develop a machine learning function for CAMARADES-NC3Rs Preclinical Systematic Review & Meta-analysis Facility (SyRF), available at app.syrf.org.uk. The facility, including the machine learning functionality, is available for public use.

## Acknowledgement

This work is supported by a grant from the Wellcome Trust & Medical Research Council (Grant Number: MR/N015665/1).

## Reference

1. Sena ES, Currie GL, McCann SK, Macleod MR, Howells DW (2014): Systematic reviews and meta-analysis of preclinical studies: why perform them and how to appraise them critically. J Cereb Blood Flow Metab; 34: 737–42

2. Elliott JH, Turner T, Clavisi O, Thomas J, Higgins JP, Mavergames C, Gruen RL (2014): Living systematic reviews: an emerging opportunity to narrow the evidence-practice gap. PLoS Med; 11: e1001603

3. Thomas, J., McNaught, J. and Ananiadou, S. (2011), Applications of text mining within systematic reviews. Res. Syn. Meth., 2: 1–14. doi:10.1002/jrsm.27

4. O’Mara-Eves, A; Thomas, J; McNaught, J; Miwa, M; Ananiadou, S; (2015) Using text mining for study identification in systematic reviews: a systematic review of current approaches. Syst Rev, 4 5-. 10.1186/2046-4053-4-5.

5. Gillian L Currie, Nicki Sherratt, Lesley A Colvin, Andrew SC Rice, Fala Cramond, Malcolm R Macleod, & Emily S Sena (2015). Search for studies using animal models of neuropathic pain [Data set]. Zenodo. http://doi.org/10.5281/zenodo.35448

6. Gillian L Currie, Nicki Sherratt, Lesley A Colvin, Andrew SC Rice, Fala Cramond, Malcolm R Macleod, & Emily S Sena (2015). Systematic review and meta-analysis of studies using animal models of neuropathic pain protocol https://drive.google.com/file/d/0B30wfjnG6aEQRS0yMVNLWmdKeDQ/view

7. Arun Varghese, Michelle Cawley, Tao Hong, Supervised clustering for automated document classification and prioritization: a case study using toxicological abstracts (2017)

8. Brian E. Howard, Jason Phillips, Kyle Miller, Arpit Tandon, Deepak Mav, Mihir R. Shah, Stephanie Holmgren, Katherine E. Pelch, Vickie Walker, Andrew A. Rooney, Malcolm Macleod, Ruchir R. Shah and Kristina Thayer (2016). SWIFT-Review: a text-mining workbench for systematic review, Syst Rev. 2016;5:87

9. Liao, Jing, (2018), shihikoo/TextMining: v1.0 (Version v1.0). Zenodo. http://doi.org/10.5281/zenodo.1194729

